# Enhanced inhibitor tolerance and increased lipid productivity through adaptive laboratory evolution in the oleaginous yeast *Metshnikowia pulcherrima*

**DOI:** 10.1101/2020.02.17.952291

**Authors:** Robert H. Hicks, Yuxin Sze, Christopher J. Chuck, Daniel A. Henk

**Affiliations:** Centre for Doctoral Training in Sustainable Chemical Technologies, Department of Biology and Biochemistry, University of Bath, Bath, BA2 7AY, United Kingdom; Department of Biology and Biochemistry, University of Bath, Bath, BA2 7AY, United Kingdom; Department of Chemical Engineering, University of Bath, Bath, BA2 7AY, United Kingdom

**Keywords:** Adaptive laboratory evolution, Microbial Oil, Lignocellulose, Fermentation Inhibitors, Bioprocessing, Biodiesel

## Abstract

Microbial lipid production from second generation feedstocks presents a sustainable route to future fuels, foods and bulk chemicals. The oleaginous yeast *Metshnikowia pulcherrima* has previously been investigated as a potential platform organism for lipid production due to its ability to be grown in non-sterile conditions and metabolising a wide range of oligo- and monosaccharide carbon sources within lignocellulosic hydrolysates. However, the generation of inhibitors from depolymerisation causes downstream bioprocessing complications, and despite *M. pulcherrima’s* comparative tolerance, their presence is deleterious to both biomass and lipid formation. Using either a single inhibitor (formic acid) or an inhibitor cocktail (formic acid, acetic acid, fufural and HMF), two strategies of adaptive laboratory evolution were performed to improve *M. pulcherrima’s* fermentation inhibitor tolerance. Using a sequential batch culturing approach, the resulting strains from both strategies had increased growth rates and reduced lag times under inhibiting conditions versus the progenitor. Interestingly, the lipid production of the inhibitor cocktail evolved strains markedly increased, with one strain producing 41% lipid by dry weight compared to 22% of the progenitor. The evolved species was cultured in a non-sterile 2L stirred tank bioreactor and accumulated lipid rapidly, yielding 6.1 g/L of lipid (35% cell dry weight) within 48 hours; a lipid productivity of 0.128 g L-1 h-1. Furthermore, the lipid profile was analogous to palm oil, consisting of 39% C16:0 and 56% C18:1 after 48 hours.

## Introduction

Microbial lipids, produced from heterotrophic organisms, are a versatile chemical feedstock that can be used to produce alternatives to fossil derived fuels and chemicals. In comparison to higher plant oils, such as palm oil, microbial oils can be produced on non-arable land, that does not compete with virgin rainforest or food production. While the lipid profile from microalgae is highly variable, generally, oleaginous yeast produce lipid profiles akin to plant oils with elevated levels of oleic and palmitic acid (Sitepu et al. 2014). Recently, we reported on the oleaginous yeast *Metschnikowia pulcherrima* that can be grown in non-sterile conditions, while having the ability to metabolise a range of oligosaccharide and monosaccharide carbon sources (Whiffin et al. 2016; Long et al. 2017; Fan et al. 2018). Despite this potential, for microbial lipids to financially compete with both fossil and vegetable oils, a low cost lignocellulosic feedstock must be used as the feedstock source.

Processing lignocellulosic feedstocks presents multiple challenges, however, one key issue is the formation of inhibitory by-products resulting from the degradation of monosaccharides produced from the cellulose and hemicellulose feedstocks. These inhibitors halt or inhibit growth and metabolism, slow down productivity and reduce overall product yields (Klinke et al. 2004). Fermentation inhibitors can be classified according to their functional group; carboxylic acid, ketone, phenolic or aldehyde (Taylor et al. 2012). Two of the most abundant are 5-hydroxymethyl furfural (HMF) and 2-furaldehyde (furfural), resulting from the dehydration of hexose or pentose sugars respectively. The most common inhibitory acids produced are formic and acetic acid, with formic acid produced as a derivative of furfural or HMF, and acetic acid produced when acetyl groups are released from hemicellulose (Radecka et al. 2015).

The inhibitory mechanism for these components is complex. HMF and furfural are non-specifically reactive with RNA, DNA and proteins as well as causing membrane damage resulting in disruptions to metabolism and cell viability (Lin et al. 2009). Acetic and formic acid can enter yeast cells via diffusion through the plasma membrane where they dissociate into acetate/formate and a proton once within the cytoplasm. Accumulation of protons leads to cytoplasm acidification causing metabolism impairment by inhibiting glycolytic enzymes and NADH dehydrogenases to increase lag times and reduce growth (Radecka et al. 2015). In addition to the individual effects of each inhibitor, they act synergistically to create negative effects that exceed the sum of each individual effect (Martín et al. 2007; Field et al. 2015).

A rational engineering approach to improve inhibitor tolerance is complex due to the vast network of molecular mechanisms involved. In addition, as genetic toolkits to accomplish such aims are unavailable within many non-model organisms, alternative phenotypic improvement strategies, such as adaptive laboratory evolution (ALE), are potentially more promising. ALE is a method whereby microorganisms are continually cultured under a selective pressure in defined conditions for periods of time to allow for the selection of advantageous phenotypes (Dragosits and Mattanovich 2013). Whilst ALE often begins with a natural isolate, it is not uncommon to first perform random mutagenesis through chemical or UV means. Many different evolution strategies have been applied to improve the biotechnological capacity of natural or genetically engineered *Saccharomyces cerevisiae* strains, for example improving tolerance to ethanol (Stanley et al. 2010) or lignocellulosic fermentation inhibitors (Wright et al. 2011) as well as improving the fermentation of non-preferred sugars such as xylose (Van Maris et al. 2007) and arabinose (Wisselink et al. 2007). There are limited examples of strain improvement through ALE within oleaginous yeast, though *Rhodococcus opacus* strains which had first undergone genetic engineering followed by mutagenesis selection were further improved by through sequentially batch culturing within increasing concentrations of lignin (Kurosawa et al. 2015).

Oleaginous yeast such as *Rhodotorula glutinis, Yarrowia lipolytica* and *Lipomyces starkeyi* tend to have a higher inhibitory tolerance compared to *S. cerevisiae* (Sitepu et al. 2014; Whiffin et al. 2016). Similarly, *Metschnikowia pulcherrima* displays a naturally high tolerance to furfural, acetic acid and HMF (Long et al. 2017). *M. pulcherrima*’s industrial potential is further improved by its inherent antimicrobial activity, allowing low cost non-sterile cultures to be performed (Santamauro et al. 2014).

The aim of this current study was to increase the fermentation inhibitor tolerance of *M. pulcherrima* whilst comparing two ALE strategies; either using a fermentation inhibitor cocktail (containing furfural, HMF, formic acid and acetic acid) as a selective pressure, or formic acid in isolation.

## Methods

### Chemicals

Unless otherwise stated, chemicals were sourced from Sigma Aldrich and used without further purification.

### Strains, strain maintenance and media

The *M. pulcherrima* strain (NCYC2580) used as the progenitor in this study was obtained from the National Collection of Yeast Cultures. Strains were maintained on malt extract agar (MEA) plates, and re-streaked on a fortnightly basis. For the preparation of overnight cultures, a single colony was inoculated into 10 mL SMB pH 5 (3% tryptic soy broth, 2.5% malt extract), and incubated at 25 °C with 200 rpm agitation. Optical densities throughout were measured at 595_nm_. Media used within ALE study was a nitrogen limited broth (NLB) consisting of: Glucose 40 g/L, (NH_4_)_2_SO_2_ 2 g/L, KH_2_PO_4_ 7 g/L, MgSO_4_ 7H_2_O 1.5 g/L, NaHPO_2_ 2 g/L and yeast extract 1 g/L. This media was autoclaved without glucose or MgSO_4_, which was added separately after autoclaving individual stock solutions. YNB medium was prepared as follows - 1.78 g/L Yeast Nitrogen Base w/o amino acids, 25 g/L glucose and variable amounts of (NH4)2SO2 per desired concentration.

### Evolution experiment

Three growth conditions were used for the initial screening experiment: NLB, NLB + 0.6 g/L formic acid and NLB + inhibitor cocktail (0.7 g/L of furfural and acetic acid, and 0.35 g/L formic acid and HMF). Triplicate overnight cultures grown in SMB were diluted to an OD ∼ 1 with PBS the following morning. Each diluted culture was used to inoculate each of the three screening conditions by adding 500 µL into 10 mL media within a glass culture tube. Cultures were incubated at 25 °C with 200 rpm agitation with OD measurements were performed at regular intervals through the 72 hour period.

For the ALE experiment, cultures were started as above using the same starting formic acid and inhibitor cocktail concentrations, with five replicate lineages for each condition. OD measurements of each culture were taken at 24 and 48 h intervals, with cultures transferred after 48 hours by inoculating 100 µL of each culture into a fresh 10 mL culture tube, taking a glycerol stock at each transfer. In the event of no growth in one or more of the culture tubes after 48 hours, successful lineages were expanded to seed new tubes to replace those lost, maintaining a total of five culture tubes per experiment. Inhibitor concentrations were increased when batches were reaching stationary phase after 48 hours to ensure that as much as practically possible exponentially growing cells were being transferred to the next batch. The evolution experiment was concluded after approximately 1000 hours where the final strains were glycerol stocked, and plated onto MEA to provide single colonies for phenotypic analysis.

### 96 well plate growth phenotypic assays

To phenotypically analyse the resulting mutant cell lines, a 96 well plate format was used. To perform these, triplicate overnight cultures for each mutant strain, as well as the progenitor were prepared as stated previously, with each diluted to an OD ∼ 1 the following morning. To prepare the 96 well plate, 140 µL of media was added to the appropriate well, and to this, 10µL of each diluted culture was added. Media blanks were plated as negative controls. Cultures were incubated within a shaking plate reader with temperature maintained at 25 °C. Optical density was read at 30 minute intervals, for a period of 48 hours unless otherwise stated within figure legends. Max growth rates were calculated using RStudio (RStudio 2015), using Ln(OD) values. Media composition and inhibitor concentrations used within the phenotypic screening assays are described within figure legends.

### Lipid extraction and profiling

Overnight cultures were prepared as previously stated, with 500 µL of the diluted overnight culture used to inoculate 10 mL of NLB or NLB supplemented with an inhibitor cocktail (concentrations used for this assay were: Acetic acid and furfural 0.7 g/L, formic acid and HMF 0.35 g/L.

The lipid extraction methodology is based upon that proposed by Bligh and Dyer (Bligh and Dyer 1959), and modified to the following:

Upon completion of culturing, 9 mL of cells were centrifuged at 13 thousand rpm, resuspended in 1 mL PBS, and centrifuged again, discarding the supernatant and freezing cell pellets instantly in liquid nitrogen. Following this, frozen cell pellets were freeze dried at -40°C for a minimum of six hours. For the cell disruption, a preweighed amount of cell pellet (ideally within the range of 10 – 100 mg) was mixed with 10 mL 6M HCl, and stirred at 80 °C for one hour. To extract lipids, 10 mL of chloroform:methanol (1:1) was added, and the mixture was stirred overnight at room temperature. To quantify the lipid weight, the lower chloroform phase was removed by hand with a glass pipette, avoiding the emulsion layer which forms between the chloroform and aqueous phase. The chloroform was then fully evaporated via rotary evaporator at 50 °C, and the remaining lipids were weighed to determine the lipid weight % of the cells.

For the lipid fatty acid profile, extracted lipid samples were transesterified with methanol and an additional 1% H_2_SO_4_, heated at 90 °C under pressure for three hours. The resulting fatty acid methyl esters were extracted with hexane, which was washed with water to remove any residual glycerol or sulfuric acid. GC-MS analysis was carried out using the Agilent 7890A Gas Chromatograph equipped with a CP-Sil capillary column (25 m x 0.250 mm internal diameter) and a He mobile phase (flow rate: 1.2 ml min^-1^), coupled with an Agilent 5975C MSD. Approximately 50 mg of each sample was dissolved in 100 ml hexane and 1 µl of each solution was loaded onto the column, pre-heated to 40 °C. This temperature was held for 1 minute and then heated to 250 °C at a rate of 10 °C min^-1^ and then held for 10 minutes. The FAME profile was calculated in reference to known standards.

### Bioreactor culturing

Overnight cultures were inoculated using a 30 mL SMB culture with an OD between 7-10 which was centrifuged and resuspended in 10 mL PBS. Culturing volume used was 1L, with 5 mL polypropylene glycol P2000 (antifoam) added upon inoculation. 80% oxygenation was maintained at the determined concentration through agitation (900 rpm max) and sparging (max 3:1 v/v), pH was maintained at 4 through the addition of nitric acid (1M) or sodium hydroxide (1M) and temperature was maintained at 20 °C. These parameters were controlled by the Fermac 320 bioreactor control unit. The cultures were performed to replicate low-cost industrial conditions, and as such, were under non-sterile conditions. Media used was NLB, or NLB supplemented with 0.7 g/L of furfural and acetic acid, and 0.35 g/L formic acid and HMF.

## Results

### NCYC2580 inhibitor tolerance and adaptive laboratory evolution

Concentrations of 0.6 g/L formic acid for the single inhibitor strategy and an inhibitor cocktail containing 0.7 g/L of furfural and acetic acid + 0.35 g/L formic acid and HMF for the multi inhibitor investigation were found to reduce 48h growth by approximately 50% compared to NLB without inhibitors, and therefore chosen as the starting concentrations for the ALE experiments (Figure 1). Inhibitor cocktail cultures displayed an extended lag time compared with the other two conditions, failing to reach an OD greater than 1 until approximately 29 hours. Despite not causing an increased lag time, the formic acid supplemented cultures instead appear exhibit biphasic growth.

**Fig. 1.**
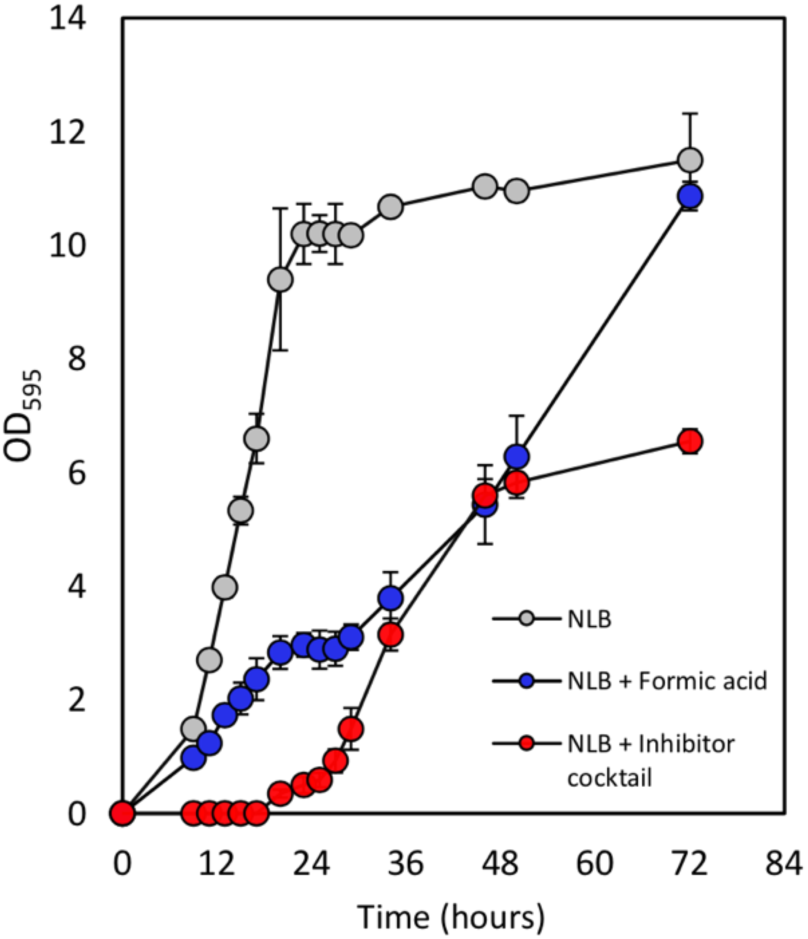
Growth profiles of NCYC2580 in the presence of inhibitors. Strains were grown in NLB as a control (grey circles), NLB with 0.6 g/L formic acid (blue circles) and NLB with an inhibitor cocktail (0.7 g/L furfural and acetic acid, and 0.35 g/L formic acid and HMF; red circles). Error bars represent the mean and standard deviation of triplicate culture

For both ALE experiments, five clonal lineages were started in parallel, and in the event of lineage loss, a successful culture was expanded to maintain a total of five cultures. Cultures were transferred after 48 h, with 100 µL of undiluted culture used to inoculate the following batch. Inhibitor concentrations were increased when the 48 hour OD indicated stationary phase was being reached, as determined by the control growth curve in Figure 1. Using this approach, the majority of cells transferred to the new batch would be growing exponentially.

### ALE: Formic acid

With a starting concentration of 0.6 g/L formic acid, OD was consistently high and the concentration across all five lineages was increased to 0.9 g/L at batch 4 (Figure 2). Increasing the formic acid concentration to 1 g/L at batch 10 resulted in reduced growth, characterised by an increased lag time (shown by the 24 h sample) in batches 10 and 11, and a decrease in the transfer OD through batches 12 to 14. All five cultures stabilised from batch 15 onwards, allowing the concentration to be increased further. Though the initial batch at 1.2 g/L formic acid saw reduced growth across all lineages, the following four batches recovered, indicating adaptation to this concentration. In total, 22 batches were performed within which a doubling of the initial formic acid concentration of 0.6 g/L to 1.2 g/L was achieved.

**Fig. 2.**
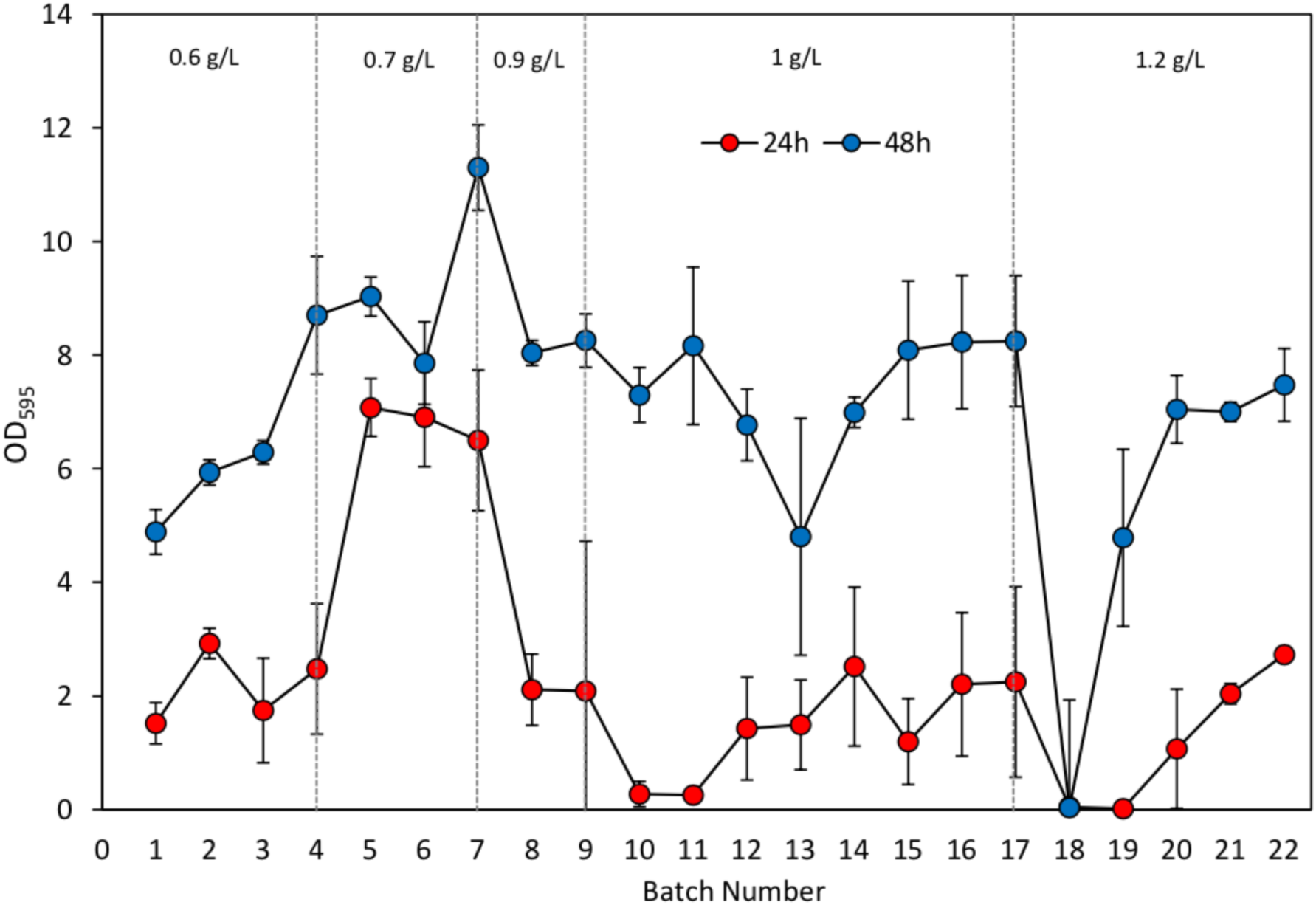
Overview of the NCYC2580 formic acid adaptive laboratory evolution experiment. Five colonies were used to inoculate five individual NLB + 0.6 g/L formic acid cultures. Cultures were transferred to fresh media after 48h growth; red and blue circles represent the OD values taken at 24 and 48 hours respectively for each batch. All five lineages continued throughout the duration of experiment, therefore error bars represent the mean and standard deviation of five cultures. The concentration of formic acid present in each batch as the experiment progresses is indicated within the plot

### ALE: Inhibitor cocktail

The second ALE strategy applied a cocktail of four inhibitors as the selective pressure, which unlike the formic acid experiment, saw frequent lineage loss within the first 9 batches (Figure 3a). In total, only one of the initial five lineages completed the entire ALE experiment, itself frequently used to expand out to five cultures. The OD data correlates with this batch to batch instability, with large fluctuations in both the 24 and 48 hour OD sample for the first 11 batches (Figure 3b). The trend onwards suggests that adaptation to these conditions; for instance, increases to 24 hour OD measurements indicated that advantageous adaptations were occurring to favour reduced lag time. These results were in correlation with that presented in Figure 3a.

**Fig. 3.**
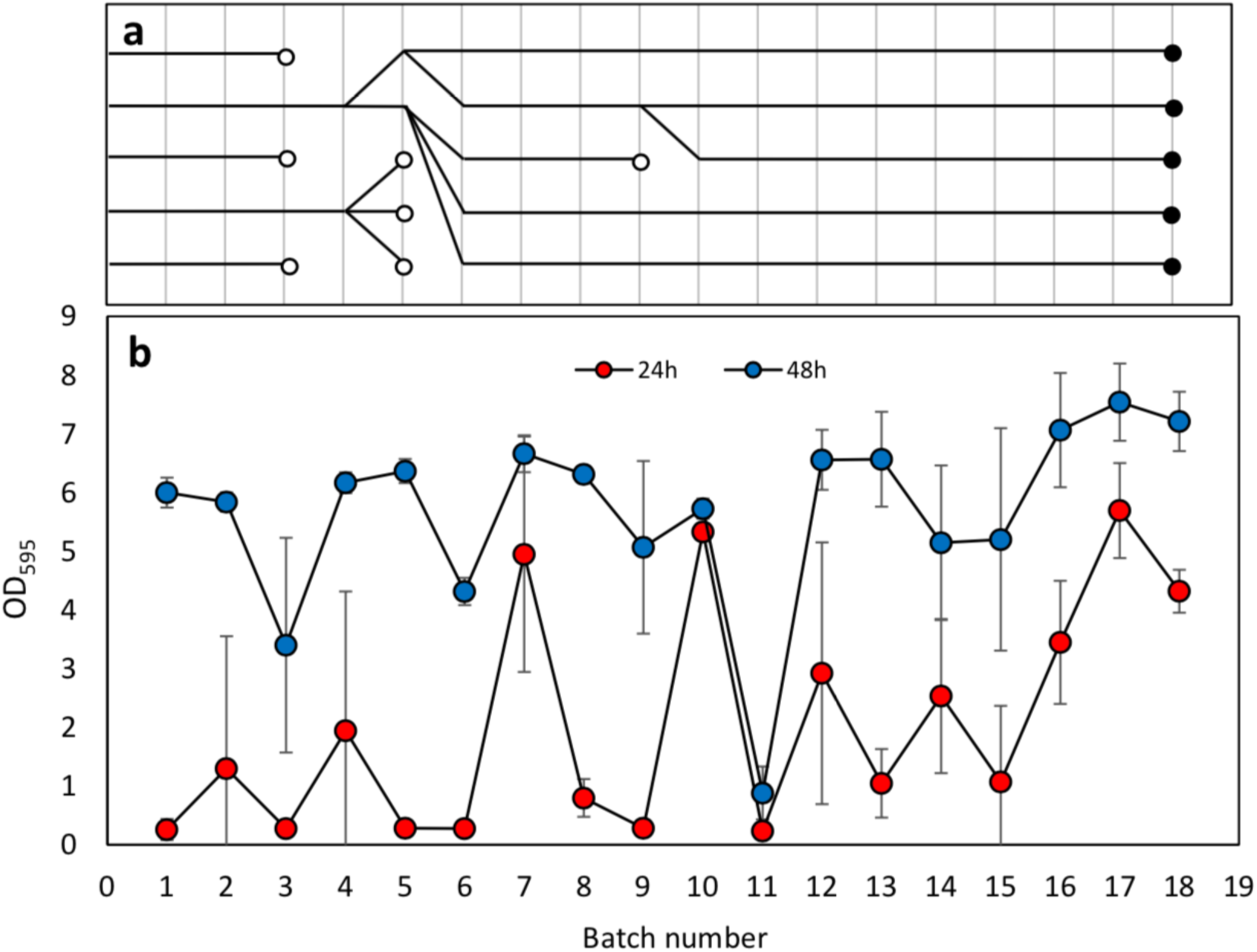
Overview of the NCYC2580 inhibitor cocktail adaptive laboratory evolution experiment. Five individual colonies were used to inoculate five NLB + inhibitor cocktail (0.7 g/L of furfural and acetic acid, and 0.35 g/L formic acid and HMF) cultures. a **-** ALE lineage tree showing successful batches replacing failed batches (clear circles) to maintain five cultures. b - Cultures were transferred to fresh media after 48h; red and blue circles represent the OD values taken at 24 and 48 hours respectively for each batch. Error bars represent the mean and standard deviation of five cultures

### Phenotypic Analysis of Evolved Strains

Upon completion of both ALE experiments, a streak plate was made from each final batch culture onto MEA agar, and a single colony from each plate was isolated onto a new MEA plate. Four out of the five lineages from each evolution strategy were taken forward for phenotypic analysis and compared to the progenitor strain. They were named as follows:

Formic acid evolved strains: F1, F2, F3, F4.

Inhibitor cocktail evolved strains: 4×1, 4×2, 4×3, 4×4.

Maximum growth rates in both rich (SMB) and defined (YNB + glucose) media determined that the evolved strains performed comparably to the progenitor when assayed in media different from the evolutionary experiments, suggesting that no discernible evolutionary trade off within these two medias had occurred (Figure 4). Strains were assayed in NLB supplemented with an inhibitor cocktail of 1 g/L acetic acid and furfural, and 0.5 g/L formic acid and HMF. In this medium, differences in the growth rate were observed, with strains evolved to the inhibitor cocktail displaying maximum growth rates double that shown by the progenitor strain (Figure 5a). The strains evolved within formic acid supplemented media also showed increased growth rates versus the progenitor, though to a lesser extent. Only the formic acid evolved strains showed increased growth rates when assayed in NLB + 0.85 g/L formic acid (Figure 5c) meaning that despite both evolution strategies yielding strains with increased growth rates when assayed with an inhibitor cocktail, here, only the formic acid evolved strains showed an increased growth rate. A decrease in the lag time across all evolved strains versus the progenitor was observed in both conditions (Figure 5b and 5d). This was most notable when grown in NLB + inhibitor cocktail where the lag time reduced by over half, with growth occurring as early as 13 hours post inoculation versus around 40 hours for the progenitor.

**Fig. 4.**
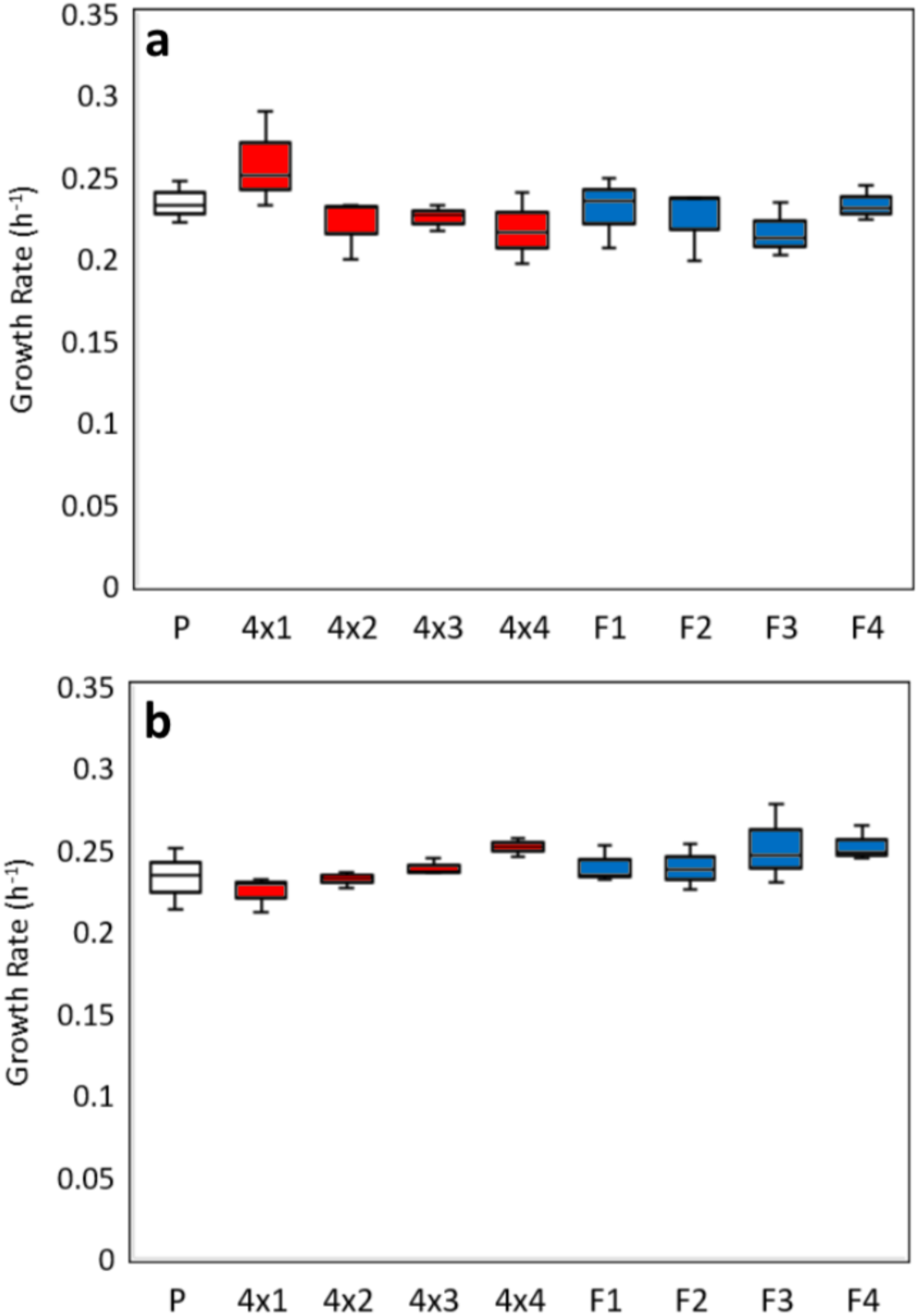
Maximum growth rates of eight mutant NYC2580 strains and the progenitor in **(a)** SMB and **(b)** YNB + glucose. Each strain was assayed in triplicate with growth performed within a 96 well plate. Max growth rates were calculated using R Studio

**Fig. 5.**
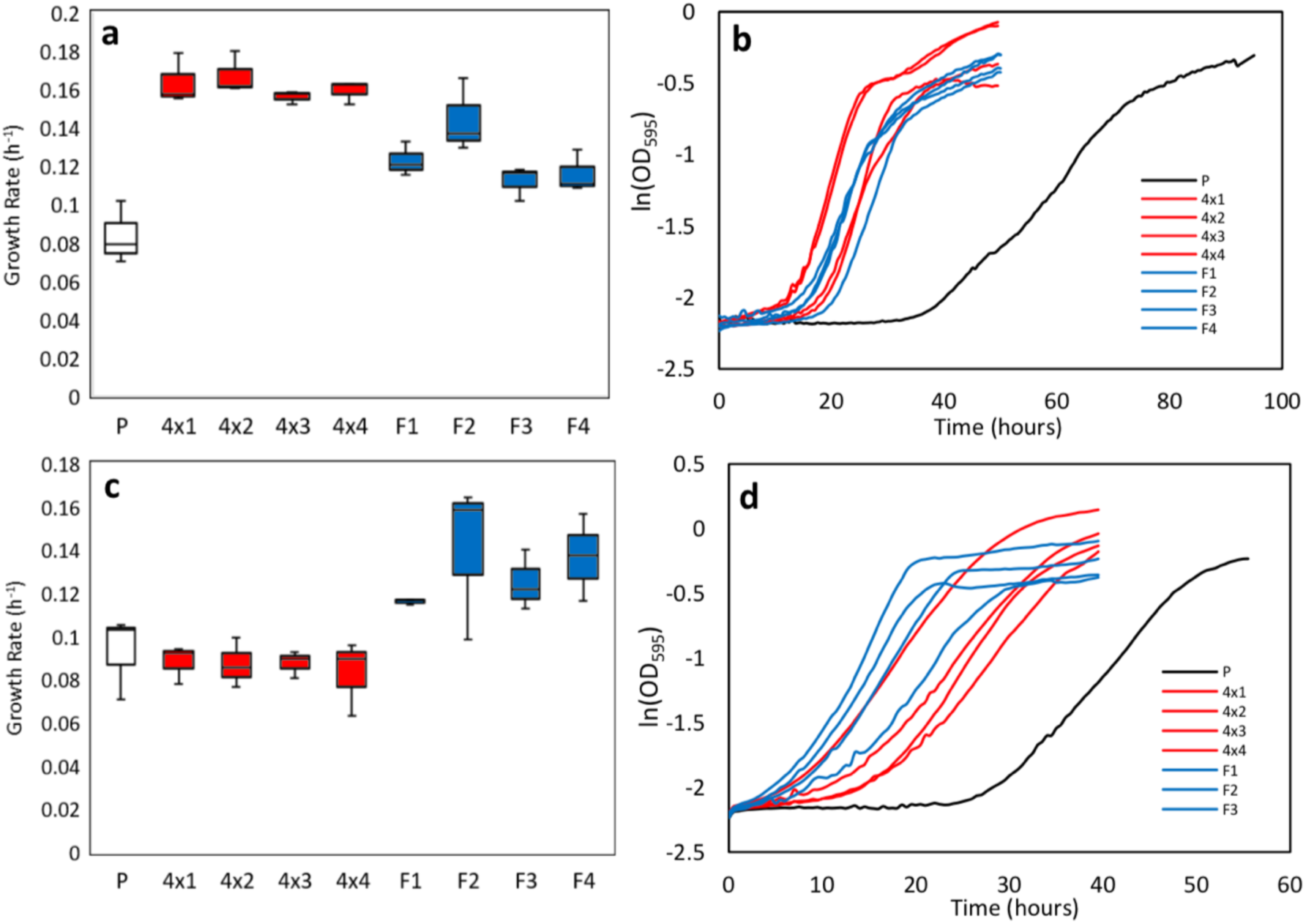
Maximum growth rates **(a)** and plotted growth rates **(b)** of eight evolved NYC2580 strains and the progenitor in NLB + inhibitor cocktail. Inhibitor cocktail consisted of: acetic acid and furfural 1 g/L, formic acid and HMF 0.5 g/L. Maximum growth rates **(c)** and plotted growth rates **(d)** of eight mutant NYC2580 strains and the progenitor cultured in NLB + 0.85 g/L formic acid. Each strain was assayed in triplicate with growth performed within a 96 well plate. Max growth rates were calculated using R Studio, curves represent the ln(OD) of each growth curve at 30 minute intervals

### Lipid production of evolved strains

Given that evolutionary trade-offs can be a significant draw back to ALE strain improvement, it is possible that the selection of inhibitor tolerant phenotypes could result in the loss of the oleaginous phenotype. To test for any effect on the lipid production, seven day cultures under inhibiting and non-inhibiting conditions were performed for the evolved strains (Figure 6). Lipid production of the progenitor strain grown in NLB without inhibitors was slightly higher (22.1%) than the four lineages evolved in formic acid (12.3-17.4%). Lipid accumulation by the four-inhibitor cocktail evolved strains increased against the progenitor to yield between 32.5 – 41% lipid by dry weight. Furthermore, despite the comparable biomass of the formic acid evolved strains and the progenitor, all inhibitor cocktail evolved strains achieved a higher biomass, with the best performing strain reaching an average of 14.4 g/L. This result was also observed when all strains were cultured within NLB + inhibitor cocktail, with the inhibitor cocktail evolved strains outperforming the others, including one mutant reaching an average biomass of 8.5 g/L with a lipid accumulation of 34.7%. The progenitor in this media had reduced biomass (4.3 g/L), but still achieved 20% lipid accumulation. In these conditions, however, the formic acid evolved strains had increased lipid production compared to when grown in control media (averaging 23% versus 15%), despite a lower biomass. Though formic acid and inhibitor cocktail evolved strains performed comparably within the growth assays, these results present strong evidence of unique adaptations occurring from each ALE strategy.

**Fig. 6.**
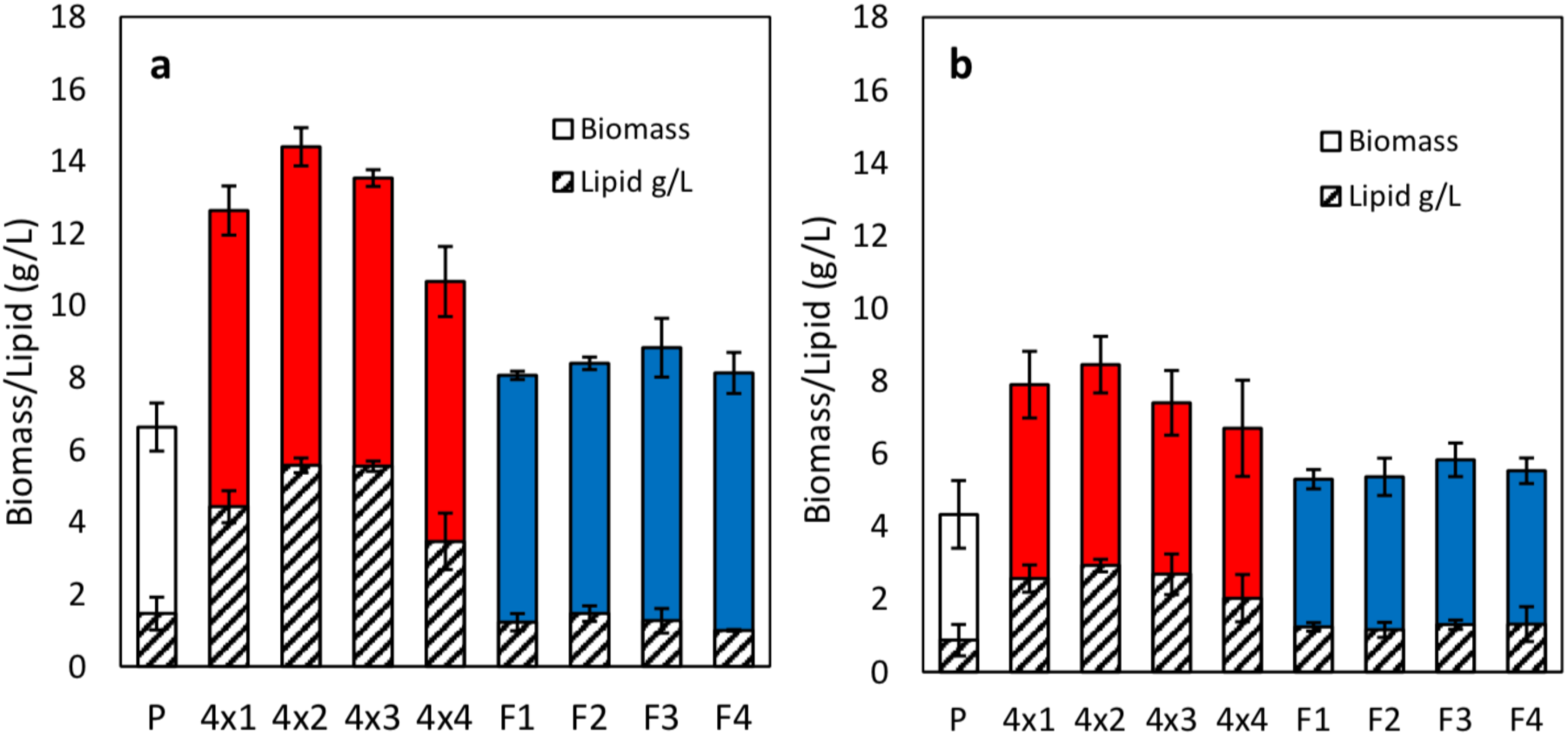
Lipid quantification and biomass of evolved strains versus the progenitor grown in NLB **(a)** and NLB + Inhibitor cocktail **(b).** Inhibitor cocktail contained acetic acid and furfural 0.7 g/L, formic acid and HMF 0.35 g/L. Triplicate samples were cultured for each strain. Error bars values represent the standard deviation of the mean

**Fig. 7.**
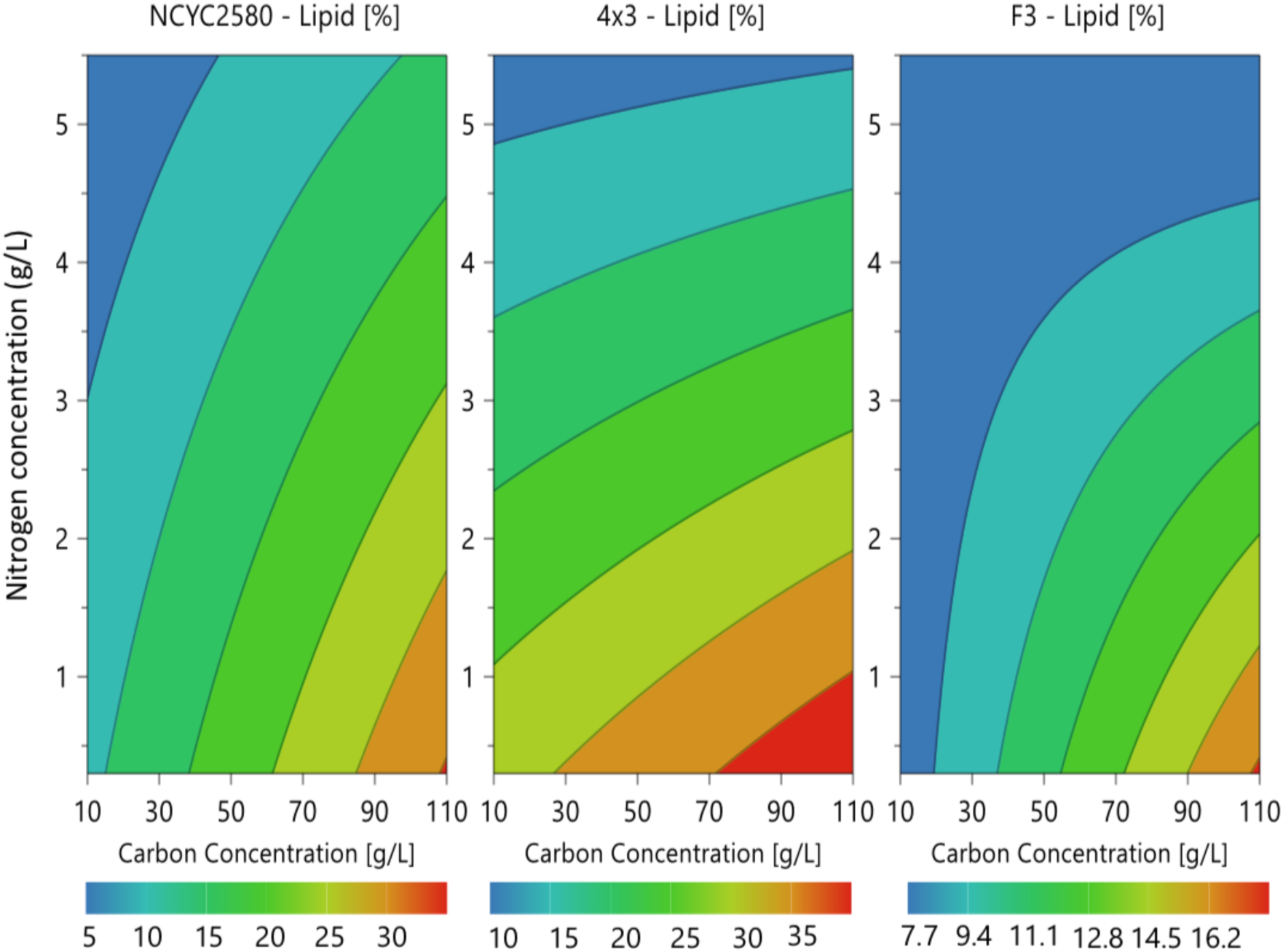
2D surface plot from a full factorial (2 levels) experimental design with the progenitor NCYC2580 and evolved strains F3 and 4×3 assayed for lipid production in response to changing concentrations of carbon and nitrogen within YNB media for seven days. Each plot represents the maximum and minimum lipid production in weight percent for the individual strain, with the range split into seven equal contours.

**Fig. 8.**
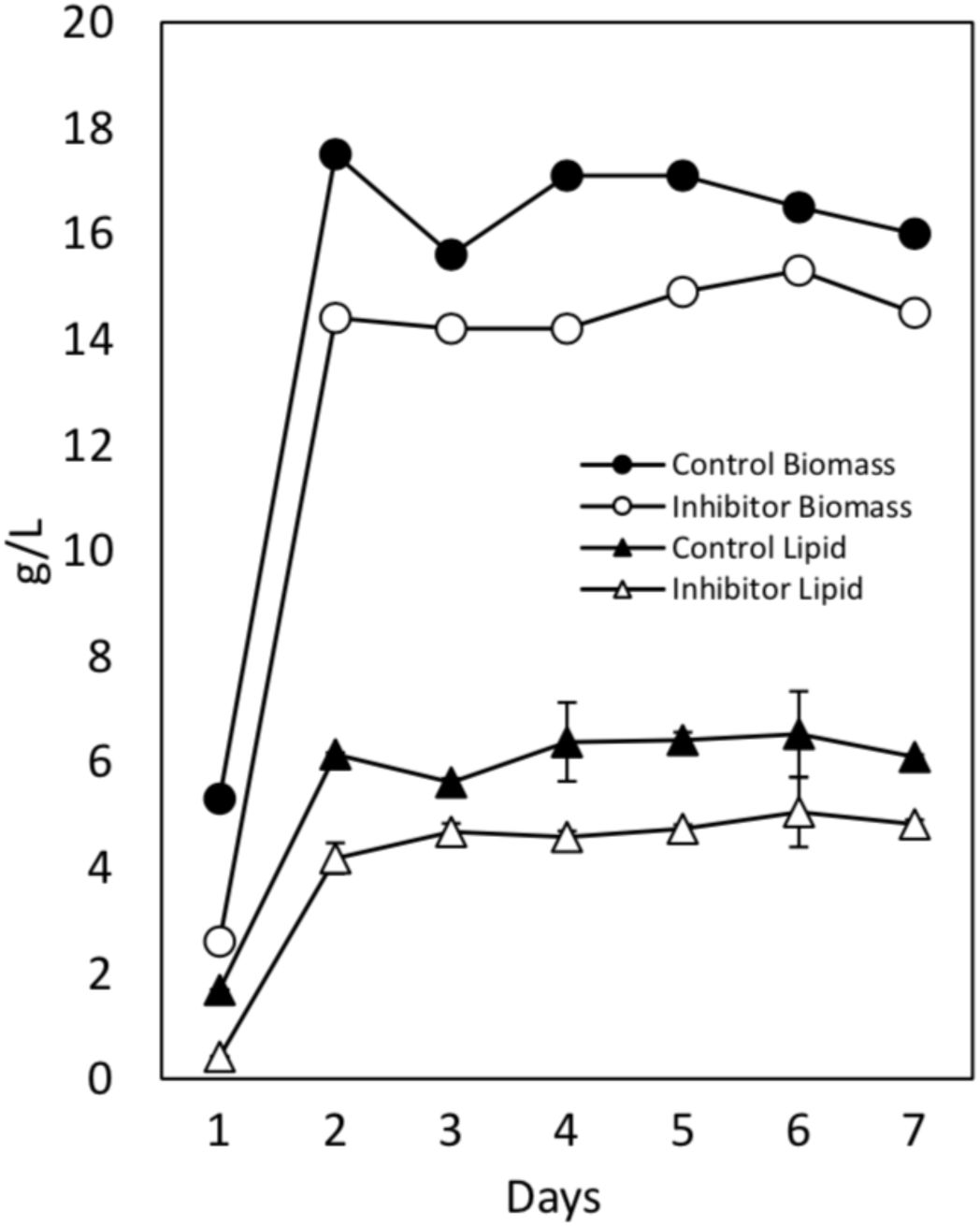
Lipid quantification and biomass of strain ‘4×3’ when grown in NLB (black circles/triangles) versus NLB + inhibitor cocktail (clear circles/triangles) under non-sterile bioreactor conditions. Inhibitor cocktail consisted of acetic acid and furfural 0.7 g/L, formic acid and HMF 0.35 g/L. Bioreactor was maintained at 20°C, pH 4 and 80% dissolved oxygen. Error bars for lipid production represent the standard deviation of triplicate samples taken at the same timepoint

Using the same samples assayed for lipid production, FAME profiling of all strains revealed further phenotypic differences. The FAME profile for the progenitor and inhibitor cocktail evolved strains varied very little when grown in the presence or absence of an inhibitor mix, though the latter had slightly increased ratios of C16:1 and C16:0 and lower amounts of C18:1 when compared to the progenitor (Table 1). The formic acid evolved strains FAME profile varied considerably to the parental however, particularly when grown without inhibitors. Ratios of C16:0 decreased to an average of 10.6% across the four strains compared to 22.5% for the progenitor, and C18:1 and C18:2 levels increased to 78.5% and 8% on average compared to 71.3% and 2.5% respectively.

**Table 1.** FAME profiles of the progenitor and evolved strains.

| NLB | C16:1 | C16:0 | C17:1 | C18:2 | C18:1 | C18:0 |
| --- | --- | --- | --- | --- | --- | --- |
| Progenitor | 3.1 | 22.5 | - | 2.5 | 71.3 | 0.5 |
| 4x1 | 4.4 | 25.6 | - | 1.2 | 67.3 | 1.5 |
| 4x2 | 8.5 | 25.9 | 0.3 | 1.6 | 61.4 | 2.3 |
| 4x3 | 7.9 | 25.3 | 0.3 | 2.5 | 62.0 | 2.0 |
| 4x4 | 3.4 | 24.2 | - | 1.4 | 68.9 | 2.1 |
| F1 | 1.2 | 11.1 | - | 5.8 | 81.2 | 0.7 |
| F2 | 2.4 | 12.3 | - | 7.2 | 77.2 | 0.8 |
| F3 | 1.2 | 8.1 | - | 11.8 | 77.6 | 1.3 |
| F4 | 3.2 | 11.0 | - | 7.2 | 78.2 | 0.5 |

| NLB + Inhibitors | C16:1 | C16:0 | C17:1 | C18:2 | C18:1 | C18:0 |
| --- | --- | --- | --- | --- | --- | --- |
| Progenitor | 1.9 | 22.0 | - | 1.8 | 73.2 | 1.1 |
| 4x1 | 5.3 | 27.5 | 0.2 | 3.5 | 62.4 | 1.1 |
| 4x2 | 4.6 | 27.0 | - | 2.7 | 63.9 | 1.8 |
| 4x3 | 4.9 | 25.7 | - | 2.1 | 65.4 | 1.8 |
| 4x4 | 3.0 | 26.1 | - | 3.4 | 66.4 | 1.1 |
| F1 | 1.9 | 17.6 | - | 1.7 | 74.9 | 3.9 |
| F2 | 2.6 | 17.1 | 0.5 | 1.7 | 75.5 | 2.7 |
| F3 | 3.0 | 16.1 | 0.7 | 1.0 | 76.7 | 2.5 |
| F4 | 3.3 | 16.1 | 0.8 | 1.1 | 76.4 | 2.3 |

NLB + inhibitor cocktail (acetic acid and furfural 0.7 g/L, formic acid and HMF 0.35 g/L). Numbers represent fatty acid composition as % of total fatty acids

### Influence of carbon-to-nitrogen ratio on lipid production in progenitor and evolved strains

Two strains; 4×3 and F3, were taken forward with the progenitor for further investigation into carbon-to-nitrogen (C:N) response and how this has been affected by each directed evolution strategy. Lipid production was assayed within a full factorial (2 levels) experimental design, with carbon and nitrogen as the input factors. The 2D surface plot models the lipid production by weight percent for each strain in response to varying C:N within the experimental space. The response range (Lipid %) for each plot ranges from the maximum and minimum lipid production values obtained for that individual strain, within the C:N ratios assayed (Figure X). All strains exhibit a similar trend in response to varying C:N ratios, accumulating the minimum amount of oil when nitrogen is high and carbon is low and maximum amounts when carbon is high and nitrogen low. The response plot for 4×3 immediately differs from F3 and the progenitor in continuing to accumulate a high amount of oil when carbon concentrations drop from their highest point, as shown by the larger red area. Another striking difference between all strains is the lipid production when both carbon and nitrogen are low (bottom left; 10 g/L glucose, 0.3 g/L ammonium sulphate). Here, F3 accumulates a low amount of oil relative to its maximum, 4×3 accumulates oil closer to its maximum and the progenitor comes in between the two. Finally, further differences are seen between the strains when comparing lipid production in top right and bottom left corners of the experimental space (high carbon, high nitrogen and low carbon, low nitrogen respectively). Here, the progenitor correlates with F3, producing a similar level of lipid at both conditions, whilst production is high for 4×3 when carbon and nitrogen concentrations are low, and decreased when concentrations are higher.

### Culturing in 2L bioreactor under non-sterile conditions

To further assess the biotechnological potential of the evolved strains, the ‘4×3’ strain evolved in the cocktail of inhibitors was taken forward for culturing in a 2L controlled bioreactor under inhibiting and non-inhibiting conditions. Here, both cultures were performed in a non-sterile manner to best represent an industrial process. Final biomass within both bioreactor conditions exceeded that observed on the smaller scale, with control media conditions achieving 16 g/L compared to 13.5 g/L within tubes, and 14.5 g/L versus 7.4 g/L with the inhibitor containing media. The biomass increase is likely due to the high oxygenation supplied within the bioreactor (80% DO) and better gas transfer. This increase in biomass also suggests that increased oxygenation aids inhibitor detoxification leading to overall higher growth. Though final biomass at day seven within the control bioreactor was higher than within culture tubes, here, peak biomass of 17.5 g/L was achieved after just 48 hours, giving a biomass productivity of 0.356 g L^-1^ h^-1^. A similar trend was also observed for lipid accumulation, where final values of 38.1% and 33.3% for control and inhibitor conditions respectively showed only a slight increase against 48 hour values of 35.1% and 29% in the tubes. This equates to a lipid productivity of 0.128 g L^-1^ h^-1^ and 0.088 g L^-1^ h^-1^ respectively.

Similarly to the smaller scale, the lipid profile did not change substantially when grown with or without inhibitors, however the level of saturated esters increased substantially compared with the tube cultures. For example, in the bioreactors the amount of C16:0 and C18:1 after 24 hours was 46% and 50% respectively against 25% and 62% when grown in the tubes (Table 2). This difference is presumably due to the increased oxygenation within the bioreactor.

**Table 2.** FAME profiles of the progenitor and evolved strains within bioreactors.

| **NLB** | **C16:1** | **C16:0** | **C18:2** | **C18:1** | **C18:0** |
| --- | --- | --- | --- | --- | --- |
| Day 1 | 1.2 | 46.0 | 0.0 | 50.8 | 1.9 |
| Day 2 | 3.2 | 38.8 | 1.3 | 55.8 | 0.9 |
| Day 3 | 4.8 | 35.2 | 2.1 | 57.4 | 0.6 |
| Day 4 | 3.9 | 37.3 | 0.0 | 57.6 | 1.2 |
| Day 5 | 3.8 | 31.5 | 1.8 | 62.1 | 0.8 |
| Day 6 | 3.8 | 30.3 | 1.9 | 62.9 | 1.1 |
| Day 7 | 3.7 | 30.3 | 1.3 | 63.8 | 0.9 |

| **NLB + Inhibitors** | **C16:1** | **C16:0** | **C18:2** | **C18:1** | **C18:0** |
| --- | --- | --- | --- | --- | --- |
| Day 1 | 0.0 | 54.9 | 0.0 | 45.1 | 0.0 |
| Day 2 | 3.8 | 39.5 | 1.9 | 54.2 | 0.6 |
| Day 3 | 4.0 | 34.5 | 1.7 | 59.0 | 0.8 |
| Day 4 | 5.4 | 31.4 | 2.4 | 59.6 | 1.2 |
| Day 5 | 5.8 | 30.0 | 2.4 | 60.6 | 1.1 |
| Day 6 | 5.2 | 30.7 | 1.9 | 60.8 | 1.3 |
| Day 7 | 3.8 | 30.6 | 1.6 | 63.3 | 0.8 |

NLB + inhibitor cocktail (acetic acid and furfural 0.7 g/L, formic acid and HMF 0.35 g/L). Numbers represent fatty acid composition as % of total fatty acids

## Discussion

### Adaptive Evolution of NCYC2580

The 48 hour batch transfer experimental design used within this study aimed to maintain cells in exponential phase as much as practically possible. In this way, stationary phase adaptation was to be minimised, and cells were exposed to a ‘fresh’ dose of inhibitors when stress tolerance is generally at its lowest (Plesset et al. 1987). The short batch time was also designed to specifically select for two industrially advantageous phenotypes; high growth rate and low lag time under inhibiting conditions, with slower growing populations eventually dropping out due to successive low frequencies at transfer. This approach however, is not favourable for selecting survival adaptations once the stationary phase was reached.

Differences between both evolution strategies were stark; inhibitor concentrations in the formic acid strategy gradually increased to 1.2 g/L from a starting point of 0.6 g/L whilst the starting concentration within the inhibitor cocktail strategy was maintained for the duration and lineages were frequently lost due to batch death. This result is most likely due to the multitude of stresses exerted by the synergistic effect of inhibitors (Martín et al. 2007). Successful ALE experiments using multi-inhibitor containing lignocellulosic hydrolysate or synthetic media containing inhibitor cocktails using a sequential batch approach are limited within the literature. Rather, studies of this kind are performed within chemostats where the maintenance of biomass leads to quicker detoxification of inhibitors by reducing the dose per cell effect (Martín et al. 2007; Smith et al. 2014). A sequential batch strategy applied by Koppram et al. in agreement with results within Figure 3**Error! Reference source not found.**, also encountered early complications, reporting a lag time increase of 35 to 50 hours within the initial batches (Koppram et al. 2012). However, as batch transfers were performed upon depletion of glucose within the media opposed to at a set time point as presented here, it is likely that this strategy is not targeted at selecting for mutants with reduced lag time or increased growth rate unlike the rationale presented here.

### Growth Analysis of Evolved Strains

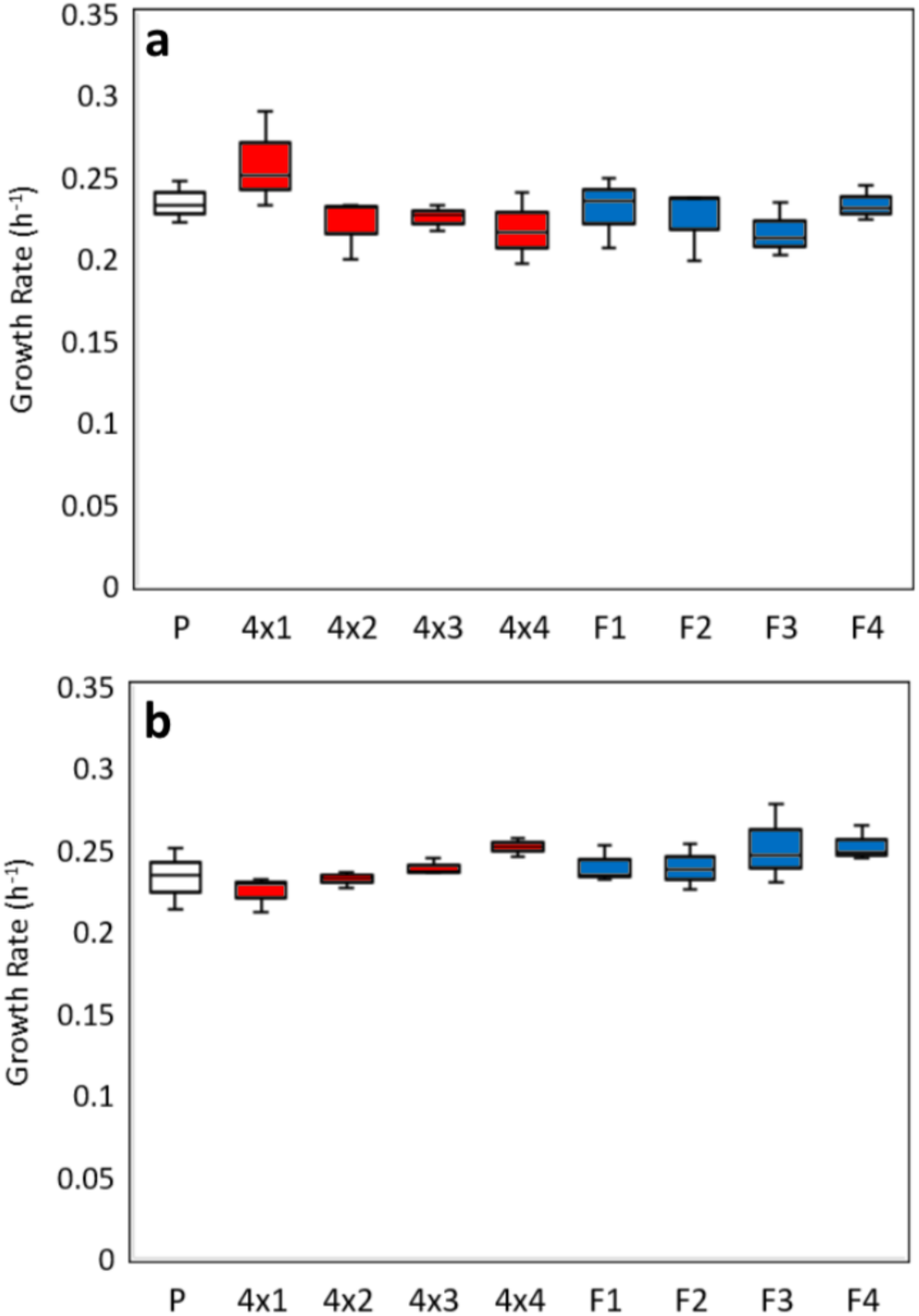

As growth rate improvements versus the progenitor were not observed within YNB + glucose and SMB (***Fig.***), it is improbable that the ALE strategies have caused advantageous mutations or copy number amplifications that increase glycolytic efficiencies leading to decreased lag time and increased growth rate. Similarly, it is unlikely that the evolved strains are faster growing due to increased glucose transportation efficiencies as has been observed within *S. cerevisiae* strains evolved to glucose-limited media (Brown et al. 1998; Dunham et al. 2002). Rather, given that growth rate improvements were only observed under inhibiting conditions, then any adaptive improvements are likely to be specific to inhibitor tolerance. Mutation or amplification candidates aiding acid tolerance include multi-drug transporters or proteins within the ABC transporter family such as Pdr12 or Azr1, the latter of which has been shown to contribute to acetic acid tolerance (Piper et al. 1998; Tenreiro et al. 2000). Increased tolerance and growth within furfural and HMF containing medias on the other hand could implicate the efficiency or abundance of furfural/HMF reducing enzymes such as Adh1, Adh6 and Sfa1 (Petersson et al. 2006). Though there are limited examples of genomic sequencing applied to strains evolved to fermentation inhibitors, Sako *et al*. found both a loss of heterozygosity within one chromosomal arm and duplication within another within an engineered *S. cerevisiae* strain evolved to inhibitor containing media, with both events involving members of the *PDR* ABC transporter gene family (Sato et al. 2014). The loss of heterozygosity event occurred within the chromosomal arm containing *PDR1*, a gene regulating the expression of multidrug resistance transporters including those associated with tolerance to gallic acid, whereas the duplication event contained *PDR18*, a gene important for tolerance to herbicides and metal ions, *PDR16*, a regulator of lipid biosynthesis and *PDR17*, a regulator of membrane composition.

As two distinct evolution strategies were performed, it was anticipated that mutants will develop adaptations specific to their own evolution condition. In this respect, as the inhibitor cocktail contained both acid and aldehyde inhibitors, it was expected that these mutants would perform well when assayed in just formic acid supplemented media which was not observed. Conversely, mutants evolved to formic acid performed strongly when assayed with an inhibitor cocktail. This is particularly surprising given the presence of HMF is known to induce long lag times in *S. cerevisiae* (Sehnem et al. 2013). These results favour the emergence of non-specific mutations to confer increased tolerance, such as the upregulation of genes such as *WHI2* encoding for a protein required to activate the general stress response and shown to increase tolerance to acetic acid and furfural both individually and in combination when overexpressed (Chen et al. 2016).

### Lipid Production of Evolved Strains

Though presence of inhibitors, whether as a single supplemented inhibitor or an inhibitor cocktail, is reported elsewhere to reduce biomass in oleaginous yeasts as shown here, their effect on lipid production varies (Chen et al. 2009; Hu et al. 2009; Yu et al. 2011). Within *Rhodosporidium toruloides* for example (Hu et al. 2009), no effect on lipid production was observed when cultured in the presence of six inhibitors, whereas the lipid accumulation of *C. curvatus* has been shown to reduce by 62% when in the presence of 1 g/L furfural (Yu et al. 2011). In this study, the lipid production of the three groups of strains were affected by inhibitors in different ways: the progenitor was unaffected, the formic acid evolved strains increased lipid production and the inhibitor cocktail evolved strains had their production reduced. Despite this, certainly the most striking result was the overall lipid production of inhibitor cocktail evolved strains increasing compared with the progenitor. Furthermore, as the formic acid evolved strains did not share this additional phenotype, it is likely that adaptation to the aldehyde inhibitors is responsible. An indication as to what adaptations may have occurred when evolving to an cocktail containing aldehyde inhibitors are given by how these compounds are processed by the cells. Within *S. cerevisiae*, furfural and HMF are both converted into less toxic compounds by NADPH-dependant enzymes (Petersson et al. 2006; Heer et al. 2009) meaning evolution to media containing both may have selected for adaptations to increase cellular concentrations of NADPH for quicker inhibitor detoxification. Mutations or gene copy number amplifications of *POS5*, responsible for the conversion of NADH to NADPH within the mitochondria, or *ALD6* which converts NADP^+^ into NADPH within the cytosol are therefore potential targets (Miyagi et al. 2009). NADPH also plays a dual role within oleaginous organisms, functioning as a critical component in fatty acid synthesis. Despite it traditionally believed that NADPH used during fatty acid synthesis was derived solely from the activity of a cytosolic malic enzyme, recent studies have revealed that NADPH produced elsewhere within the cell can also be used to this end. These include the activities of glucose-6-phosphate dehydrogenase (G6PD) or 6-phosphogluconate dehydrogenase (PGD) in the pentose phosphate pathway, meaning that several alternative NADPH sources could be involved in the increased lipogenesis observed by these strains (Zhang et al. 2007; Chen et al. 2015). Given this, it is possible therefore that if a mutation has caused cellular levels of NADPH to be greater than that of the progenitor, particularly in conditions where inhibitors are not present, that fatty acid synthesis is providing a ‘sink’ for the excess NADPH. Furthermore, if these mutations were combined with a gene duplication to the lipid biosynthesis regulator *PDR16* as was observed within a *S. cerevisiae* strain evolved to fermentation inhibitors (Sato et al. 2014), then their synergistic effect may be responsible for this new phenotype.

An alternate explanation for increased lipid production within the multi inhibitor evolved strains could be due to an altered C:N response. Applying an experimental design to assay C:N response of two evolved strains, 4×3 and F3, revealed distinct differences from its progenitor stage. In particular, high lipid accumulation by 4×3 was shown to occur over a broader range of C:N ratios assayed compared to the progenitor, with the reverse true for strain F3. As both strategies took place within nitrogen limited media, evolution to display a nitrogen limited, and therefore lipid rich, phenotype at lower ratios of carbon to nitrogen does not seem to have occurred due to the dissimilar C:N response by these two evolved strains. Again therefore it suggests that adaptation to the inhibitors themselves, rather than adaptation to nitrogen limited media, is responsible for the altered C:N response of 4×3. Indeed, it is reported that genes indicating nitrogen starvation were up regulated in *S. cerevisiae* in response to HMF and furfural supplemented media, with it further stated that NADPH flux usually directed towards ammonium assimilation under control conditions was instead redirected to detoxification of HMF and furfural (Ask et al. 2013).

Although overall lipid production was influenced by the presence of inhibitors (Figure 6), the FAME profile for all strains remained relatively unchanged (Table 1), agreeing with work by Hu et al. and Yu et al. (Hu et al. 2009; Yu et al. 2011). What did affect the FAME profile however was culture, age as shown within the non-sterile bioreactor cultures (Table 2). Under both inhibiting and non-inhibiting conditions, initially high levels of C16:0 within the first 48 hours of growth reduced and C18:1 and C16:1 levels increased as the cultures progressed. Here, it is likely that C16:0 fatty acids are either being converted enzymatically to C16:1 through the activity of desaturases such as *OLE1* (Ledesma-Amaro and Nicaud 2016), or further elongated (either before or after desaturation to C16:1) to C18:1. Maintaining a more saturated lipid, suitable as a palm oil substitute, would therefore require the cessation of culturing after 48 hours, or targeting enzymes such as *OLE1* for deletion. Whereas if a more monounsaturated product was required, for biodiesel production, then the yeast would need a longer time in the bioreactor.

The strain presented in this study appears unique amongst oleaginous organisms in producing a microbial oil high in C16:0, with *Cryptococcus curvatu* (28%), *R. glutinis* (18%) and *Y. lipolytica* (11%) having a notably lower proportion (Ageitos et al. 2011). The narrow FAME profile of this organism, consisting almost exclusively of C16:0 and C18:1 fatty acids, also appears contrary to other oleaginous yeast. The FAME profile of *Y. lipolytica* and *Trichosporon pullulans* for example is reported to consist 51% and 24% of C18:2 respectively, whilst *C. curvatus* and *R. graminis* produce 15% and 12% C18:0; FAME’s *M. pulcherrima* appears to produce only in trace amounts (Beopoulos et al. 2009; Ageitos et al. 2011).

This study emphasises the effectiveness of adaptive evolution to generate improved phenotypes in instances where genetic engineering is either not available, or, as with the aims of this study, difficult to approach with rational design due to the vast network of molecular mechanisms. By employing different strategies, though both successful with respect to increasing inhibitor tolerance, adaptive evolution can provide strain improvements beyond the initial aim.

## Acknowledgments

This research has been funded by the Industrial Biotechnology Catalyst (Innovate UK, BBSRC, EPSRC) to support the translation, development and commercialisation of innovative Industrial Biotechnology processes (EP/N013522/1) and by the EPSRC through the Centre for Doctoral Training in Sustainable Chemical Technologies (EP/L016354/1).

## Author Contributions

ALE was planned by RHH and performed by YS. RHH planned and conducted all further work, and wrote the manuscript. CJC and DAH advised the study, and revised the manuscript. All authors read and approved the final manuscript.

